# The Conceptualization and Preliminary Evaluation of a Dynamic, Mechanistic Mathematical Model to Assess the Water Footprint of Beef Cattle Production

**DOI:** 10.1101/2020.04.14.028324

**Authors:** Hector M. Menendez, Alberto S. Atzori, Luis O. Tedeschi

## Abstract

The water footprint assessment method has helped to bring livestock water use to the forefront of research to address water challenges under the ecological footprint perspective. The current assessment methods of water use make a meaningful assessment of livestock water use difficult as they are mainly static, thus poorly adaptable to understand future scenarios of water use and requirements. They lack the integration of fundamental ruminant nutrition and growth equations within a dynamic context that accounts for short and long-term behavior and time delays associated with economically important beef producing areas. This study utilized the System Dynamics methodology to conceptualize a water footprint for ruminants within a dynamic and mechanistic modeling framework. The problem of beef cattle livestock water footprint assessment was articulated, and a dynamic hypothesis was formed to represent the Texas livestock water use system as the initial step in developing the Texas Beef Water Footprint model (**TXWFB**). The fulfillment of the dynamic hypothesis required the development of three causal loop diagrams (**CLD**): cattle population, growth and nutrition, and the livestock water footprint. The CLD provided a framework that captured the daily water footprint of beef (**WF**_**B**_) of the cow-calf, stocker, and feedlot phases and the entire beef supply chain. Preliminary simulations captured the oscillatory behavior of the Texas cattle population and overshoot and collapse behavior, under conditions when regional livestock water resources became scarce. Sensitivity analysis from the hypothesized CLD structures indicated that forage quality was less of an impact on the daily WF_B_ of each cattle phase compared to the use of high concentrate feeds. This study provided a framework concept for the development of a dynamic water footprint model for Texan’s beef cattle production and water sustainability.

## Introduction

Global demand for water resources has created much pressure for sectors that have large water footprints (1). These sectors include industry, household, and agriculture. Within the agriculture sector, livestock production has received a large amount of scrutiny. It has created the impetus for assessing and reducing livestock water use. Initial efforts to understand, quantify, and standardize livestock water use have been made by the Water Footprint Network, the International Standards Organization, and the Food and Agriculture Organization, amongst many other methods (2). Most livestock water use methods include some form of the water footprint assessment (**WFA**) method developed by Hoekstra in 2002 (3). The WFA includes the quantification of three specific water types, green (rainfed), blue (ground or surface), and grey (waste treatment) water uses that account for the total direct, and indirect water (i.e., virtual water) used to produce a product (4).

Current livestock water use literature indicates that beef cattle have the most significant water use among different livestock, making the quantification of beef water use the predominant area of interest and investigation. However, beef cattle also have the broadest range of water footprint values caused by numerous water use methodologies and the associated interpretations of their results combined with broad differences in production efficiency. The most notable difference between beef water use methodologies is how they account for green, blue, and grey waters, if at all, and their functional unit. Consistency is another problem. For example, livestock water use may be in liters of water per kg of live weight, hundredweight, carcass weight, or boneless beef. Additionally, regional and environmental considerations may further change the functional unit into an index of scarcity based on available and returned water use over a given period (e.g., a month) (5).

Legesse et al. (2) provide a comprehensive review of current methodologies to assess beef livestock water use, and more recently, the Food and Agriculture Organization (6) published a standardized methodology for livestock water use systems and supply chains. Standardizing the evaluation of livestock water use is essential to determine the actual resource consumption and allocation per area (e.g., country or state). Water use evaluation also helps to indicate levels of unsustainable water use and water scarcity and provides a benchmark to improve upon (7–9). Therefore, one can optimize the management of world freshwater resources based on the water footprint, because it considers direct and indirect use of all components of the water usage geographically (e.g., country, province, state) and temporally. Despite these tremendous efforts, the current methodologies are still based on static frameworks, limited periods of assessment, and neglect to capture the structure of the problem of livestock water use quantification. It presents a big limit to forecast water use by production sectors, to formulate or simulate possibles scenarios for future water management in the production areas and to evaluate technical strategies to improve water use efficiency in the production chains. Animals are grown in dynamic, continuous systems that receive feedback from climate, soil, available feed, and supply chain dynamics. Tedeschi and Fox (10) described the evolution of computer-based simulation models to adequately evaluate ruminant livestock feed and water requirements. However, there has been no attempt to link animal growth, the plane of nutrition, and water requirement in a dynamic, long-term assessment. Current livestock water use methodologies are missing these equations and fail to account for continuous diurnal physiological and environmental processes (Fig 1). Furthermore, these processes are often delayed in time and are dynamically interconnected through powerful feedback mechanisms that influence daily and total water use. Turner et al. (11), for example, described numerous agriculture examples of the need for a dynamic methodology to solve complex agriculture challenges more adequately. Few researchers (12–18) have published dynamic models relating to cattle that provide meaningful perspective and high-leverage policies to influence systems over the long-term. Therefore, there is a need to more critically evaluate the beef water footprint (**WF**_**B**_) using available cutting-edge ruminant nutrition and growth equations (10) with a dynamic framework to advance available water footprint assessment methodologies and perform various, highly functional and rapid, policy analyses.

**Fig 1.**
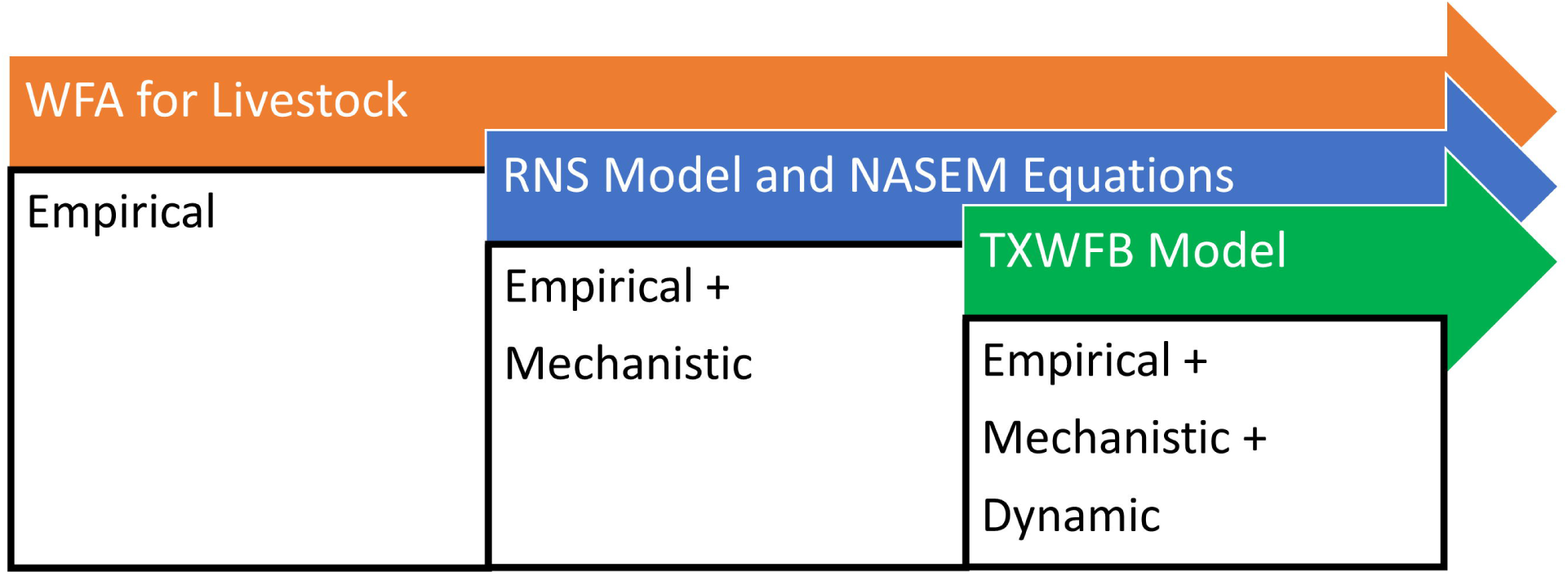
Overview of equations types in various models. Equation sources include the Water Footprint Assessment (**WFA**), Ruminant Nutrition System (**RNS**), National Academies of Science, Engineering and Medicine (**NASEM**) and the Texas Beef Cattle Water Footprint (**TXWFB**) Models and their equation types.

Within the United States, an area that is economically dependent on beef cattle production is Texas, one of the top-five cattle producing states (19). Texas also covers many large and diverse geographical and climatic regions in which the three major phases of beef production exists. Additionally, management of cattle in each region have their own respective ecological and resource limitations (e.g. drought, unproductive soils) and benefits (water availability, favorable climates). Thus, this study focuses on Texas, at a state level of aggregation, for the development the conceptual framework due to its robust representation of beef cattle production under many ecological and climatic regions.

The first objective of this study was to describe relevant information about the water use processes involved in beef cattle production at the ranching level for developing a dynamic water footprint model. The second objective was to conceptualize these processes into a dynamic framework (i.e., dynamic hypothesis) that would allow a dynamic water footprint model to be developed and evaluated. The third objective was to provide preliminary calibration of the proposed dynamic framework to improve its ability to capture important feedback signals in cattle water use over time and assess its suitability for model formulation (i.e., the addition of equations).

## Material and methods

### System dynamics methodology

The primary objective of improving existing WF_B_ WFA was accomplished by applying the System Dynamics (**SD**) methodology in developing our model framework. The SD methodology is an approach well adapted to understanding complex systems (20,21). Complexity in water management is related to non-linear dynamics which are non-linear relationships between variables that may have a tremendous influence on a system’s behavior or none. For example, cattle may accumulate heat over a given time. However, it is not until a point at which their body can no longer maintain homeostasis that declines in feed intake are observed. These non-linear dynamics are guided by feedback mechanisms (i.e., loops) that can be reinforcing (e.g., accumulation of heat) or balancing (e.g., decrease in feed intake) whose influences are delayed over time. Often, the unintended consequences of decisions in the cattle industry are difficult to understand and change because managers react to events (e.g., diminished feed intake) instead of understanding the structure (i.e., environmental exposure, body condition, breed type, feed type) that is driving the system.

This approach uses a high level of aggregation to describe the overarching structure of a system in which the problem exists and captures important non-linear dynamics, feedbacks, and time delays that are responsible for the inertia that drives the system behavior. The modeling process is based on well defined steps approaching system understanding. Step one enables the problem to be clearly articulated and defined, explicitly states what the model aims to understand and resolves the purpose of the model. Step two, dynamic hypothesis formulation, describes the model boundaries clearly, it consists of a concise statement or causal loop diagram that clearly describes the hypothesized endogenous (variables that are part of the feedback dynamics) structure that drives the problem behavior. Exogenous variables, according to SD nomenclature, are variables that do not receive feedback from variables within the system. Problem articulation and formulation of a dynamic hypothesis are iterative processes and should improve throughout the modeling process. This study utilized steps one and two of the SD method to articulate the problem and develop several causal loop diagrams (**CLD**; diagram of endogenous variables) that resulted in a dynamic hypothesis CLD and statement. Steps one and two were accomplished by conducting an extensive review of literature, identifying key variables, using expert knowledge, and published SD models to best capture the structure of the system in which the problem exists.

### Problem statement and dynamic hypothesis

The dynamic problem statement included the definition of the lack of knowledge limiting an accurate determination of Texas beef waterfootprint and its use to formulate, and test, adequate policies for efficient resource use. The problem statement focused on the model conceptualization with the inclusion of key variables from the beef chain (regarding herd dynamics, productivity, management of growing phases, commercial trade, environmental constraints, etc) and strongly affecting water use at farm and regional level.

The dynamic hypothesis expresses the hypothesized structure of the system in which the problem of current WFA WF_B_ assessment exists (Fig 2) and was used to develop specific conceptual submodels that, when aggregated, represent the core Texas Beef Water Footprint model (TXWFB) parameters and boundaries. Local, domestic, and international water resources attribute to the total input of green, blue, and grey waters used for Texas beef cattle production. Forage and crop production yield (t/ha) and water use (m^3^/ha) efficiency affect the specific water demand (m^3^/t) of pasture and feedstuffs and depend upon the management practices of multiple beef cattle stakeholders: ranchers, landowners, hay suppliers, farmers, feed mills, and feedlots. Stakeholder management practices also alter the water demands for cooling, chemical mixing, cleaning, waste treatment, dust control, nutrient and drinking requirements, and animal growth and performance (i.e., obtaining mature weight, size, and carcass quality; Fig 2: loops R1, R2, R3). The centralization and decentralization of the three major cattle production phases (i.e., cow-calf, stocker, and feedlot) across a wide range of climate and environmental conditions, sub-tropical to arid, create a disparity in water used at each phase and collectively between all phases; resulting from significant gaps in communication across the supply chain (Fig 2: loops R4, B1). Communication gaps include the lack of knowledge transfer of cattle water use levels as they progress across the supply chain.

**Fig 2.**
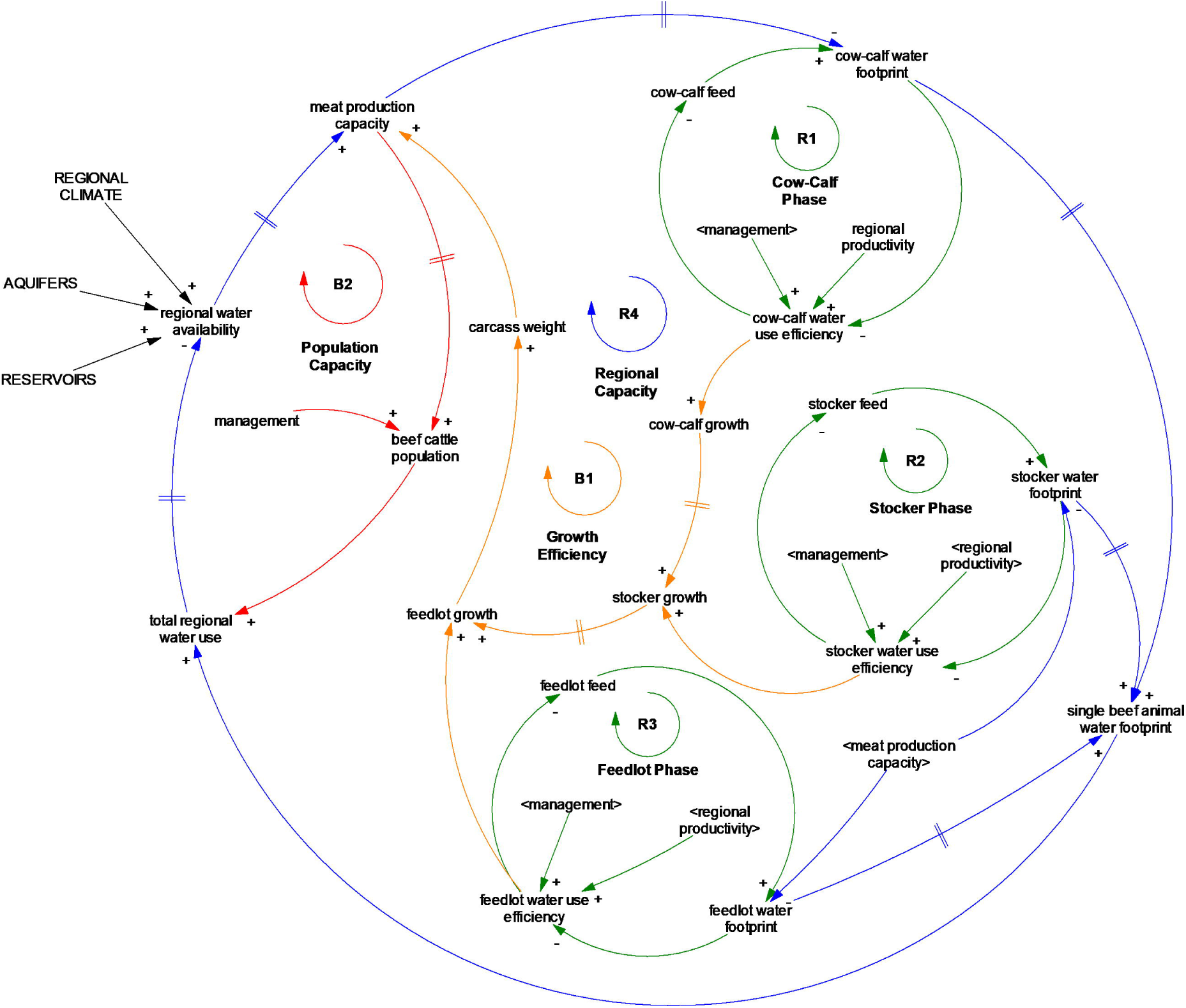
The dynamic hypothesis causal loop diagram. Blue arrows indicate linkages between variables and plus (+) and minus (-) signs denote variables relationship (i.e., same or opposite directionality). Perpendicular lines on linkages indicate time delays for processes to occur between variables. The circular arrows indicate the direction of the feedback pathway (i.e., clockwise or counterclockwise) where the R is reinforcing and the B is balancing. Bolded wording indicates the name of each loop.

Further, smaller operations tend to be more heterogeneous in management and water use efficiency but still account for a large proportion of total water use, while more extensive operations are mainly homogenous in terms of management. Therefore, the variation of the Texas beef cattle population, water use, existing resource availability, and limitations impact water allocation (m^3^), water cost (USD/m^3^), total meat production (kg), and marginal profitability ($/kg meat; Fig 2: loop B2). Local, state, and federal agencies may incentivize or disincentivize beef cattle stakeholders’ level of water use efficiency for beef cattle operations as water scarcity or consumer perception change. Identification of actual water use and comparative advantage amongst Texas regions and feedstuff type/production efficiency [i.e., specific water use(m^3^/t crop or forage)] provides baseline TXWFB measurements for sustainable water use and helps to bridge communication gaps between major beef cattle production phases for high leverage, water-reducing, improvements. Providing a baseline TXWFB value to show current and marked improvement in beef water use may also relieve consumer’s perception that consuming beef is unsustainable and harmful to the environment. Instead, TXWFB has the potential to indicate to consumers that areas dominated by grasslands and rangelands have the appropriate ecological capacity to produce beef and that alternatives such as increased grain crop production (lower-water costs) in lieu of beef may have unintended economic (decrease of US competitiveness) and environmental consequences (land and water degradation, the loss of nutrient cycling, wildlife/insect habitat and ecological goods and services).

### Model conceptualization

A professional version of dynamic modeling software, Vensim DSS, was used to visualize three specific submodels in the form of CLD from the factors identified in the dynamic hypothesis statement. The conceptual submodels include (1) cattle population, (2) growth and nutrition, and (3) the livestock water footprint. The methods section references Figs 2-5 and uses panel A (the top half) to visualize the model structure. In the results section Figs with two panels (A and B) are describe independtly and together to convey the resulting behavior of each conceptual structure.

**Fig 3.**
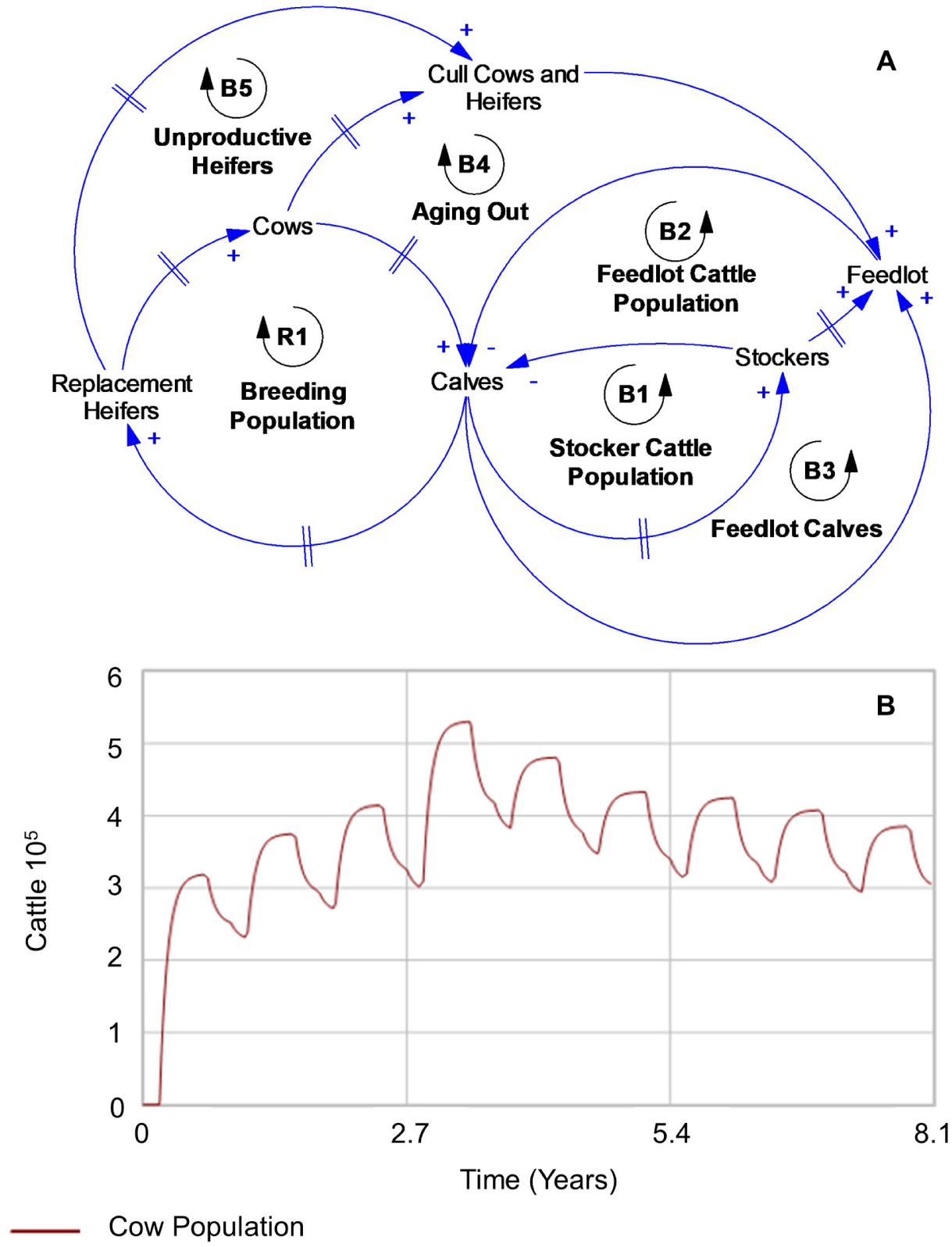
The dynamic structure of the beef cattle population (A) and an example of oscillatory behavior from the structure of the cattle population system (B). Blue arrows indicate linkages between variables and plus (+) and minus (-) signs denote variables relationship (i.e., same or opposite directionality). Perpendicular lines on linkages indicate time delays for processes to occur between variables. The circular arrows indicate the direction of the feedback pathway (i.e., clockwise or counterclockwise) where the R is reinforcing and the B is balancing. Bolded wording indicates the name of each loop.

**Fig 4.**
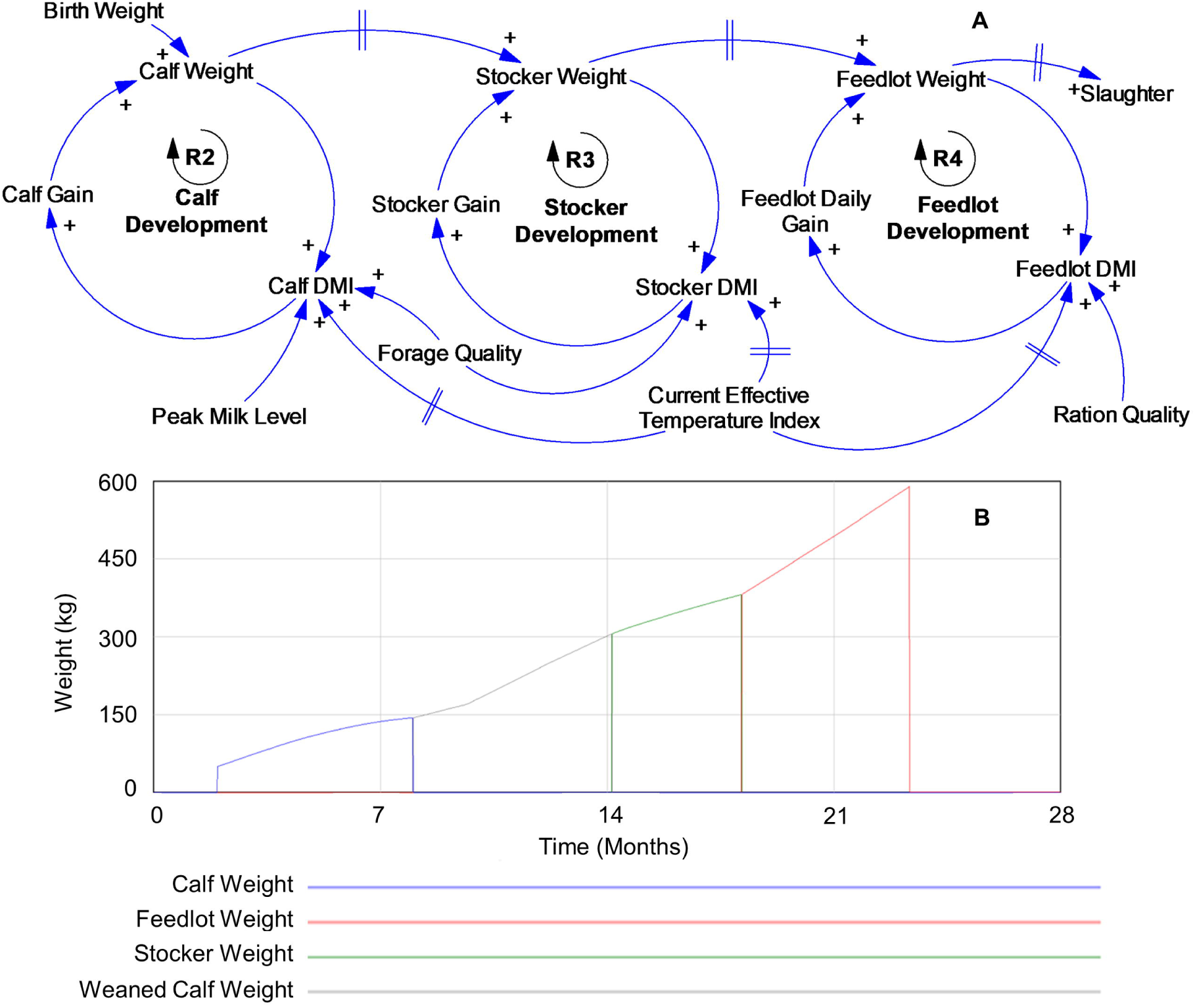
The dynamic structure of cow-calf, stocker, and feedlot growth and nutrition and interlinkages across the supply chain (A) and an example of cattle growth (kg/day) behavior (B). Where DMI means dry matter intake. Blue arrows indicate linkages between variables and plus (+) and minus (-) signs denote variables relationship (i.e., same or opposite directionality). Perpendicular lines on linkages indicate time delays for processes to occur between variables. The circular arrows indicate the direction of the feedback pathway (i.e., clockwise or counterclockwise) where the R is reinforcing and the B is balancing. Bolded wording indicates the name of each loop. The weaned calf and stocker weight (panel B) represent the entire stocker stage after weaning.

**Fig 5.**
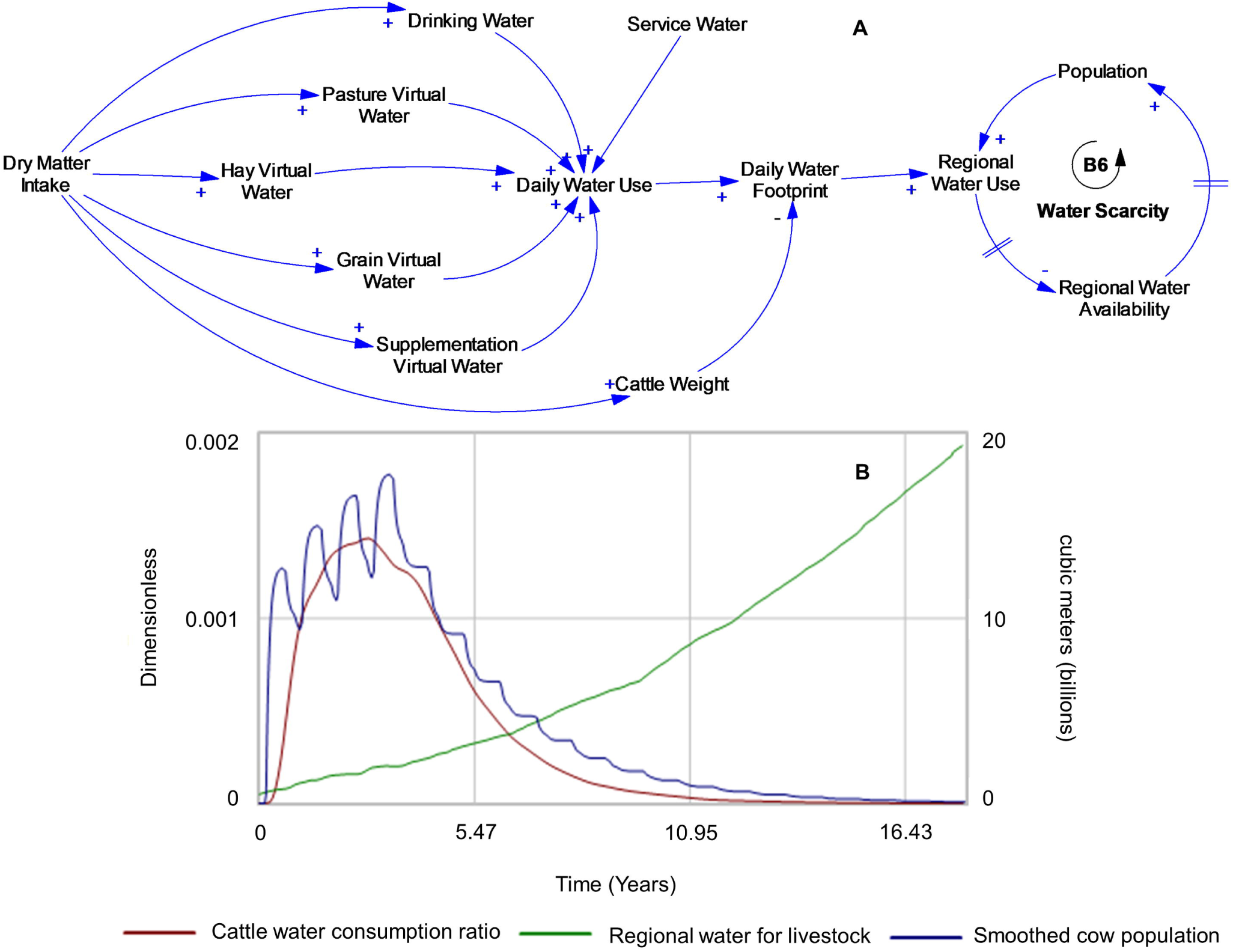
The daily livestock water footprint CLD (A) and an example of preliminary TXWFB model simulation of overshoot and collapse behavior (B). Blue arrows indicate linkages between variables and plus (+) and minus (-) signs denote variables relationship (i.e., same or opposite directionality). Perpendicular lines on linkages indicate time delays for processes to occur between variables. The circular arrows indicate the direction of the feedback pathway (i.e., clockwise or counterclockwise) where the R is reinforcing and the B is balancing. Bolded wording indicates the name of each loop. The daily cattle water consumption is a ratio between the daily cattle water use (m^3^/d) and the total available water for livestock (m^3^). The smoothed cow population is the averaged daily mature cow (cattle, not shown on-axis).

### Cattle population

The production of beef cattle in the United States is segmented (cow-calf, stocker, and feedlot) and follow the same general pattern like many regions around the world (10), but specific intricacies exist among regions, even within the United States, that impact the WF_B_ (Fig 3). Cattle management decisions for reproduction, growth, and sales are influenced by available resources (i.e., time, finances, feed, water), and economics. Collectively, short and long-term decisions influence cattle populations throughout the cow-calf, stocker, and feedlot phases. First, the cow-calf phase (Fig 3A) serves as the primary reinforcing structure that ensures beef cattle will be available each year through the development of replacement heifers and maintenance of a mature cow herd (Fig 3A: loop R1 Breeding Population).

Replacement heifers, after a two-year delay, will return to the mature cow herd and contribute to the next generation of progeny. This is a closed-loop system, meaning that the feedback exists between the number of calves born and the number of replacement animals available to sustain a commercially viable population. Consequently, mature cows, calves, and replacement heifers within this feedback loop are consuming resources. Calves not selected for re-breeding (heifers or steers) enter the portion of the beef cattle supply chain that terminates at slaughter when a desired mature weight is obtained. The desired number of stocker and feedlot cattle reduce the calves that are available for rebreeding (Fig 3A: loops B1 Stocker Population; B2 Feedlot Cattle Population). The duration of resource allocation to cattle varies greatly throughout the beef supply chain. For example, weaned calves may remain at the same ranch and region, or they may be sold and shipped to an entirely different region when entering a new phase (e.g., stocker or feedlot phases). Calves may also be sold directly to a feedlot phase and circumvent the stocker phase (i.e., Fig 3A; loop B3, Feedlot Calves). Within the cow-calf phase, some cattle fail to be productive and do not or cannot produce calves (Fig 3A: loops B4 Aging Out; B5 Unproductive Heifers) and are culled for meat production which decreases (balancing action) the total breeding population. Similar to the population loop in Fig 3A., stocker and feedlot cattle are consuming resources for different durations and at different water use intensities; some regions may have higher or lower water use intensities associated with forages, grains, and from climatic conditions. Overall, Fig 3A provides the fundamental structure of the primary reinforcing mechanism (R1: Breeding Population) and balancing mechanisms that sustain the beef cattle population and maintains a stable supply of beef for consumption. Similar structures have been used for hogs (22) and beef cattle (13,23). Conceptualization of this part of the problem, daily WF_B_ assessment, captures the importance of resource use duration and variation that exists within and across the beef cattle supply chain.

### Growth and nutrition

Population dynamics (Fig 3A) drive the behavior of the system and influence the nutrition and growth dynamics within and across each major cattle production phase (Fig 4); cow-calf, stocker, feedlot (Fig 4A). Each phase contains a reinforcing feedback mechanism that influences weight (kg). Weight drives the amount of dry matter intake (DMI), which influences the rate of gain (kg/day). Suckling calves (not weaned) consume milk primarily and then shift to forage-based diets as they mature (Fig 3A: loop R1 Breeding Population, Fig 4A: loop R2 Calf Development) (24).

Maturity influences physiological and anatomical characteristics determining feed inputs and virtual water use. Upon weaning of suckling calves, cessation of milk production from cows (dams), the calves enter the stocker stage and consume primarily forages. Forage quality and duration of the stocker phase influences the rate of growth and weight that stocker cattle will obtain during this phase (Fig 4A: loop R3 Stocker Development). Milk and forage inputs generally utilize less virtual water (m^3^) as the cow-calf and stocker phases obtain resources from pasture, grasslands, and rangelands. The majority of stocker cattle will progress to the feedlot where their diet is transitioned over a three-week period from forages (pasture or roughages) to a high concentrate-based ration; total mixed ration (**TMR;** Fig 4A: loop R4 Feedlot Development). High concentrate TMRs result in higher rates of weight (muscle and fat; kg/day) deposition during this period, known as the finishing phase. However, the water use associated with high concentrate diets [e.g., virtual water of grains (m^3^/t)] is much higher than most pasture and rangeland inputs (25). Aside from diet type, environmental factors are very influential to cattle growth and performance regardless of the cattle phase. Extreme temperatures have been shown to influence the DMI of cattle, and this exogenous (non-feedback variable) has been described by Tedeschi and Fox (10) as the current effective temperature index (CETI). Although the actual equation is not described, the structure of the CETI equation is important as it is not an instantaneous calculation; it was developed to account for the impact of climate on animals over a period (usually 15 days). The CETI captures the physical delay of heat accumulation and dissipation in cattle affecting the animal’s daily DMI and drinking water intake. In complex systems, delays, such as the delay captured using the CETI, allows the hypothesized daily water footprint model structure to account for short and long-term environmental impacts on cattle nutrition. The cow-calf, stocker, and feedlot phases (Fig 4A: loops R2, R3, R4) share the same nutrition and growth structure, indicating that if the adequate quality of nutrients is available and environmental conditions do not limit feed, then the cattle in each phase will continue to gain weight and increase DMI and their daily gain. Additionally, the three development loops (Fig 4A: loops R2; R3; R4) are connected between each phase as the animal progresses across the beef cattle supply chain (Fig 4A). Interestingly, the growth and nutritional dynamics are managed at each production phase (i.e., independent). However, the stocker and feedlot phases are influenced by the initial weight of the cattle they are receiving, which impacts their nutritional requirements, potential growth, and the duration required to achieve desired weight to reach slaughter (i.e., dependent; supply chain dynamics). Ultimately feed, and growth dynamics affect daily cattle water use and the daily water footprint at each phase (cow-calf, stocker, feedlot) and aggregated water use across the beef cattle supply chain (Fig 4A).

### Livestock water footprint

The dry matter intake of the cattle population is the main driver of the cattle performances (weight gain) and also of the water footprint. The daily water footprint is an aggregation of drinking water and service water consumption (direct water use), and also of pasture, hay, supplementation, and concentrates (e.g., grains) water uses (virtual water) that represent the daily water use required to achieve cattle growth (26–28). In this study the virtual water of feeds is determined using the specific water demand (m^3^/t) approach calculates is the amount of water (green or blue; m^3^/ha; i.e., evapotranspiration) used to produce a given amount of forage or grains (t/ha) (27,29). The daily water footprint inputs are quantitatively dependent on the amount of feed intake and are connected to the growth and nutrition feedback dynamics for each cattle phase (i.e., cow-calf, stocker, and feedlot, see growth and nutrition section above and Fig 4A). The daily water use (L/d) is then divided by the daily weight gain of boneless beef (kg/d) to obtain a daily water footprint (L/kg). Daily boneless beef is the percent of boneless beef of the live animal weight gain at a given physiological stage (i.e., young to mature; nutrient demands). The, daily water footprint, differs from the WFA for livestock in that the WFA only reports a WF_B_ at slaughter versus a continuous value reported by the TXWFB.

Determining the daily water footprint reflects the average for cattle in similar beef production supply chains. However, it does not reflect the total resource use or its impact. Assessing beef cattle resource use and impact requires that the average WF_B_ be multiplied by the cattle population in a given region (Fig 5; Fig 5A). As the cattle population increases, so do the quantity of regional water use, and this reinforcing relationship is accelerated or slowed by the instantaneous daily WF_B_. If the regional beef water use exceeds available water for livestock, then resources (e.g., drinking water, forages, and grain) may become scarce or exceed the operating budget to maintain profitability or, in extreme cases, cattle may die from lack of feed or water. Therefore, the cattle population (Fig 3A) receives a balancing feedback action until regional water levels are sufficient to sustain a given cattle population (Fig 5A: loop B6 Water Scarcity). The balancing loop of Fig 5A points out the carrying capacity of the system which is represented by the water availability and the sustainability of this resource. The fast or exponential growth of the beef sector will generate high pressure on the regional water use, especially enhancing feed production and crop cultivations, with increasing demand for blue man-managed water. A region will probably support further beef cattle population growth until the delayed, and unintended effect of water scarcity collapses the population. The dynamics of overshoots and collapses are well known in natural resource exploitation (30).

### Sensitivity analysis

Upon completion of the population, growth and nutrition, and water footprint CLD the model was parameterized with coefficients and equations from the Ruminant Nutrition System (**RNS**) (10) and the Nutrient Requirements for Beef Cattle (31). This preliminary calibration was then used to perform preliminary behavioral tests and sensitivity analyses on critical components of the calibrated model. Behavior tests are the evaluation of simulation results of a variable of interest to identify its pattern of behavior over time (e.g., reinforcing, balancing, oscillation). Three behavior tests were performed. The first test evaluated the behavior of the mature cow population by simulating the population submodel for ∼8 years. The second test evaluated the typical behavior of the cattle growth and nutrition submodel across the cow-calf, stocker, and feedlot phases. The third test evaluated the mature cow population, regional water availability, and the proportion of daily cattle water consumption for ∼18 years in a water-limited scenario. Sensitivity analysis is a quantitative and qualitative model test that indicates the amount of variation of a variable of interest (e.g., daily water footprint) from the alteration of a constant variable [e.g., total digestible nutrients (**TDN**)]. The first sensitivity analysis varied TDN values of forage (pasture and hay) and grains (i.e., individual components that comprise the TMR). The second sensitivity analysis ran 1000 simulations of the same TDN values as sensitivity test one (±10%) impact on the daily WF_B_. The third sensitivity analysis evaluated the daily WF_B_ by altering the daily dry matter (**DM**) forage production rates 25, 75, 100 (kg/ha/day) in each cattle phase to adjust the specific water demand (SWD; m^3^/t) of forages and varied the SWD of grain crops (corn = 30 to 1500, soybean = 1500 to 5000, and distillers grain = 0 to 1500 m^3^/t) (32). Collectively, the CLD, behavioral, and sensitivity methods provide preliminary results that support the overarching hypothesized dynamic structure required for the TXWFB to model a daily WF_B_.

## Results and discussion

The problem of the limitations of current WFA methodologies has been articulated in the introduction section, and steps one and two of the SD method were used to articulate the problem and form a dynamic hypothesis. The results of this study include the major CLD diagrams, their associated behavior, and sensitivity to parameters changes of important variables that influence the WF_B_.

### Causal loop structures and behaviors

The cattle population CLD identified one dominant reinforcing loop of the Texas cattle population (Fig 3A: loop R1). Five unique balancing loops were identified that cause the cattle population to decrease (Fig 3A: loops B1-5). The preliminary simulation results of these six loops indicated that the Texas population has an oscillatory behavior (Fig 3B). When synchronized with cattle population dynamics, three reinforcing loops were identified for the cattle growth and nutrition CLD (Fig 4A) and resulted in the reinforcing growth behavior of cattle weight for each cattle phase (Fig 4B). The growth and nutrition dynamics directly influenced the water consumption of the beef supply chain and the regional resource use (Fig 5A). Further, the dynamic structure of the growth and nutrition CLD resulted in a positive linkage (i.e., if one increases or decreases the linked variable does so as well) between DMI and four daily virtual (i.e., indirect) water uses (Fig 5A). Drinking and service water were the two direct water uses identified in the WF_B_ CLD (Fig 5A). The WF_B_ CLD resulted in the linkage of daily cattle weight (i.e., boneless beef) and total daily cattle water use, which represents the daily water footprint (Fig 5A). Regional water scarcity and population dynamics were also developed in the WF_B_ CLD and resulted in one balancing loop to account for cattle population carrying capacity (Fig 5A: loop B6 Water Scarcity). Preliminary simulation of the water scarcity loop was able to create overshoot and collapse behavior (Fig 5B). Preliminary simulation of these cattle growth and nutrition and water footprint dynamics indicated that non-linear relationships and delays play a role in daily weight gain relative to the daily WF_B_ (Fig 6; Fig 6AB).

**Fig 6.**
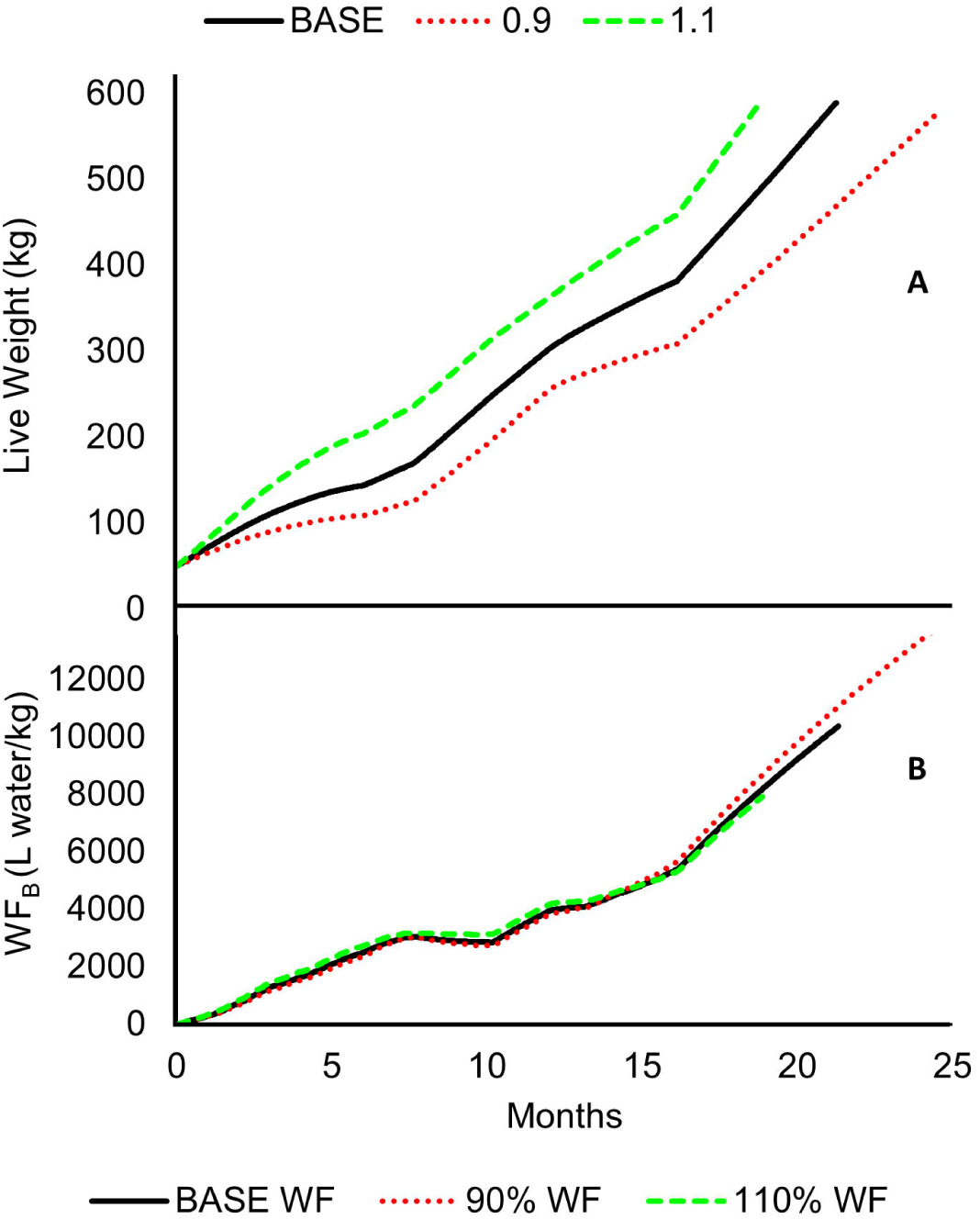
Preliminary growth and nutrition sensitivity analysis. The three scenarios include base (640 days; 21.3 months), 90% of total digestible nutrient values (**TDN**; 749 days; 24.9 months), and 110% TDN nutrition (566 days; 18.6 months) impact on weight (A) and the daily WF_B_ (B).

### Sensitivity analysis

Initial sensitivity analyses were performed to evaluate changes in cattle live weight (kg/d) and the daily WF_B_ (L water/kg boneless beef). The first sensitivity analysis of TDN, three scenarios, indicated that the time required to reach the desired mature weight (589 kg live weight) and the daily WF_B_ were increased and decreased with 90% and 110% baseline TDN values, respectively (Fig 6AB). The second sensitivity analysis of TDN, 1000 scenarios, showed that the feedlot stage had the most extensive daily WF_B_ variability compared to the cow-calf and stocker stages (Fig 7; Fig 7A). The results of the third sensitivity analysis of forage and crop SWD indicated that production efficiency had a major impact on the daily WF_B_ across all beef cattle phases and that the daily WF_B_ can be higher in cow-calf or stocker phases and lower in the feedlot phase in some circumstances (Fig 7B).

**Fig 7.**
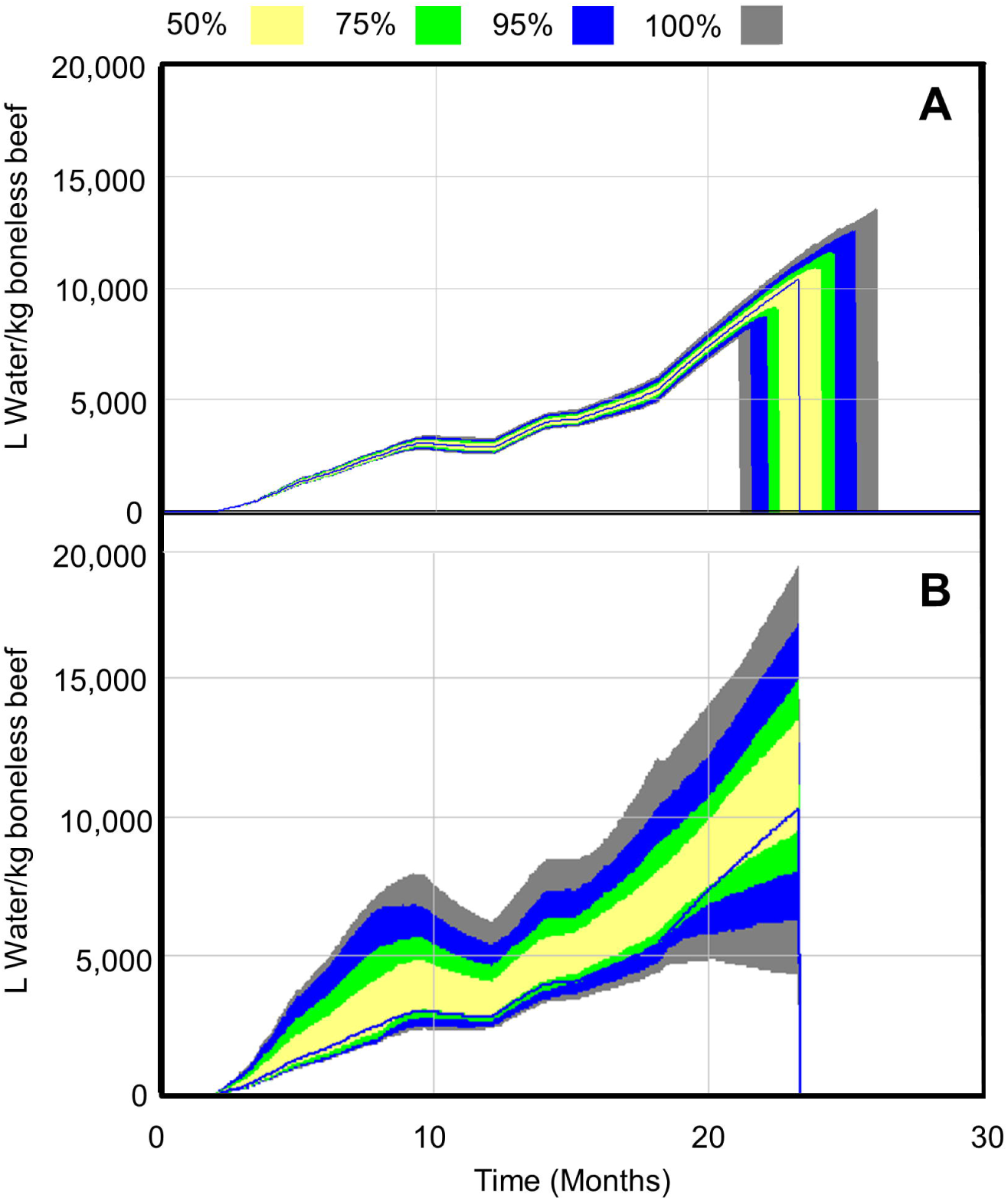
Preliminary sensitivity analysis. Preliminary sensitivity analysis of ±10% of TDN for forages and grains across all beef cattle phases (i.e., cow-calf, stocker, feedlot) on the daily water footprint (L water/kg boneless beef; A). Preliminary sensitivity analysis of annual dry matter forage production, low (6,838 kg/ha), medium (20,510 kg/ha) to high (27,350 kg/ha) on the daily beef water footprint (L water/kg boneless beef; WF) and SWD of grains (B).Yellow, green, blue, and grey colors represent the percent of simulated water footprint values within given ranges.

## Discussion of the dynamic framework

The SD methodology was successfully employed and contextualized existing WFA methods into a dynamic conceptual framework. The hypothesized structure from the dynamic hypothesis led to the conceptualization and preliminary behavioral and sensitivity analysis of cattle population, growth and nutrition, and water footprint dynamics. The population model produced the oscillatory behavior seen in other existing animal population models (Fig 3B) (20,30,33). The behavior of the growth and nutrition models was also as expected to show an increase in weight-dependent upon nutrient quality, environment, and management (Figs 3A and 5A) (10,31,34,35). Similarly, behavioral tests of the daily WF_B_ produced the expected increase of WF_B_ levels as the time required to reach slaughter was prolonged, especially in the feedlot stage (Figs 3A and 5A), and these behaviors have been identified by Mekkonen and Hoekstra in 2012 (36). A combination of the cattle population and the daily cattle water use relative to regionally available water, in a water-limited scenario, resulted in the expected overshoot and collapse behavior. The overshoot and collapse behavior is a harsh system response that reflects unsustainable cattle population growth and large fluctuations in beef production and price (i.e., supply and demand). Sterman (20) gives examples of overshoot and collapse in dynamic models and their market and business strategy implications. Understanding and avoiding this pitfall is critical for beef cattle stakeholders as livestock water use limitations, and pressure for more sustainable beef production grow. Behavioral results increased the confidence that the WF_B_ framework was adequate for livestock WFA and identified the long-term behavior types (e.g., oscillation, exponential growth/decay) of cattle population, growth and nutrition, and WF_B_ within this system.

Agreement of CLD behavior with existing models and publications led to the sensitivity analysis of critical constant parameters that were thought to be influential to several variables of interest, the daily WF_B_, cattle population, livestock water availability, and cattle weight. Sensitivity analysis of TDN revealed expected sensitivity that cattle weight and daily WF_B_ are generally lower when TDN is low, which is consistent with ruminant nutrition and growth principles (Fig 6AB) (31). However, increased simulations (1,000) revealed that TDN had the most significant impact at the feedlot stage from TMRs with high concentrate diets, and these results are reasonable as cattle put on a substantial proportion of their weight and increase DMI during this phase. This aligns well with existing WF_B_ WFA that emphasizes the large water cost of feedlots (see studies mentioned in the introduction section above). Interestingly, the sensitivity analysis of forage and crop SWD revealed that the feedlot stage might have a lower daily WF_B_ than the cow-calf or stocker stages (Fig 7AB). Ambiguity exists for SWD values as pasture and hay (forage) growth, even within a region, depends on the climate, management of stocking rates, and soil fertility of the land. Thus, the improvement of forage water use efficiencies is likely a high-leverage solution to improve the WF_B_, and this should be investigated. Furthermore, the estimation of SWD values for crops is even more confusing than forage production. The current WFA method assesses high-resolution spatial areas to determine the SWD of grain crops in the same region as the cattle. However, in Texas, feed resources are procured from domestic (United States) and international sources, i.e., it is unlikely that grains consumed by cattle came from the same region. Therefore, the sensitivity analysis of grain crop SWD (e.g., corn and soybeans) that accounts for national SWD variation provides a more robust analysis of actual green and blue virtual water uses and their impact on the daily WF_B_.

### Sustainability of production phases

Often highly connected production supply chains lack communication between phases or segments that cause unintended consequences that are delayed in time, making it challenging to identify the root cause of a problem (e.g., WF_B_). The dynamic framework in the current study provides a means for dialogue about the identified causal feedback mechanisms and delays that drive the WF_B_ within and across the cow-calf, stocker, and feedlot phases. For example, Fig 8, couples the existing feedback structures in this system and extends them to potential considerations for sustainability, production efficiency, marketing of beef products, and consumer consensus as the public becomes more aware of livestock water use intensities and the associated levels of sustainability. The balancing loop of the water scarcity embodies the sustainability of the beef sector connected to resource uses (Figs 4A: loop B6). A primary goal should be to keep the daily water footprint of beef production as low as possible. Increases in the daily water footprint of the agricultural sectors will decrease the regional water availability, depress the growth of food production, and water use efficiency (Fig 8: loop R6 Cattle Growth).

**Fig 8.**
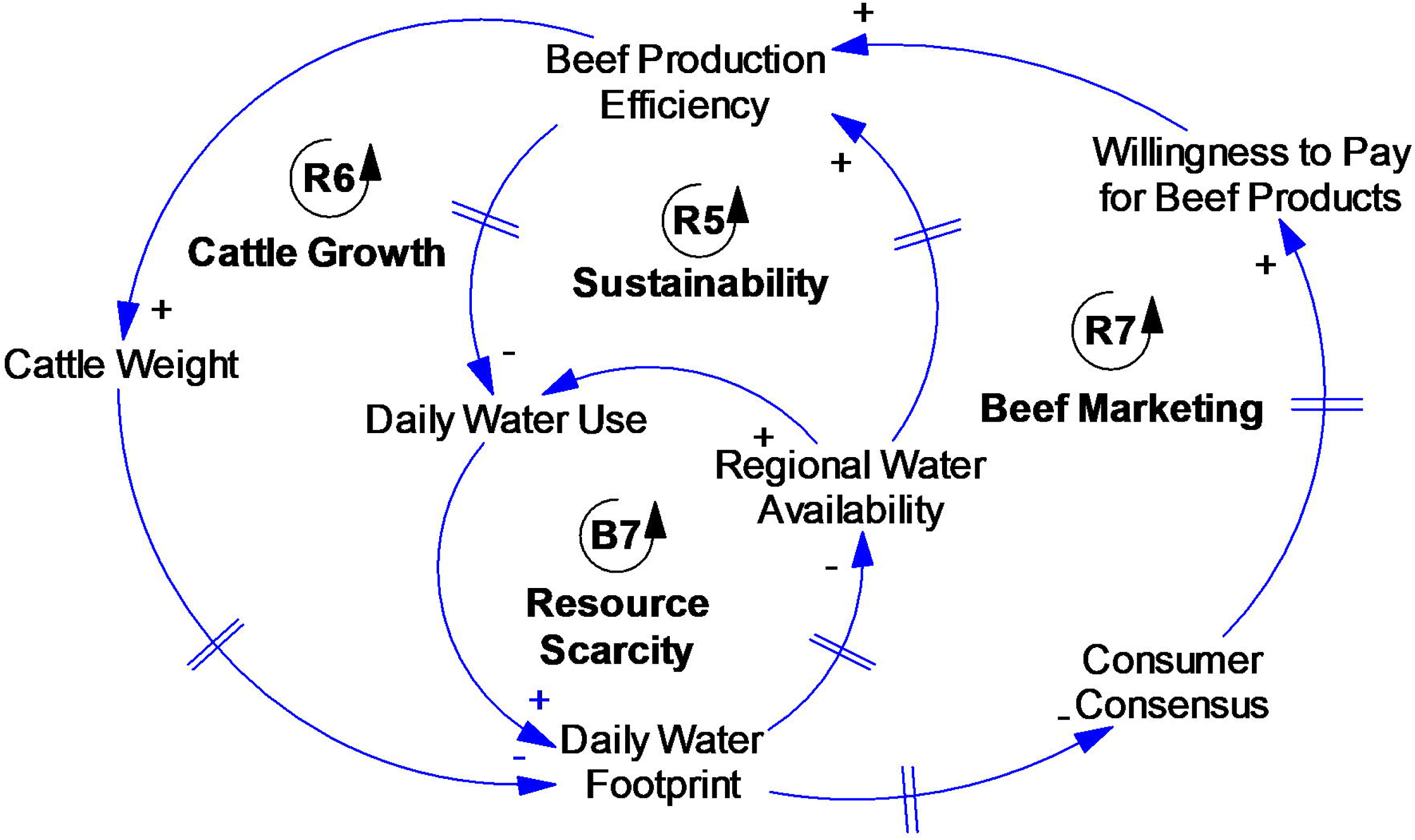
Causal loop diagram of the critical dynamic structure (described in results) and the anticipated feedback structure of public perception, marketing, and efficiency on the long-term beef water footprint of the beef livestock supply chain.

It will also reduce the food supply as a direct consequence. One of the most powerful leverage points to reach high sustainability levels could be to reduce the daily water footprint and keep increasing production efficiency (37). Improving efficiency will increase cattle weight per unit of consumed water, or reduce the daily water use, ending in a lower daily water footprint. Additionally, improvements in efficiency will allow water resources to be spared and provide greater availability of regional water, which will make the beef sector grow (Fig 8: loop B7 Resource Scarcity). As a positive side effect, the lower environmental impact of the WF_B_ might increase consumer consensus for the beef sector and the willingness to pay for environmentally friendly products. Increased sales from environmentally friendly beef products will keep the pressure to continue to improve cattle production efficiency for low impact beef (Fig 8: loop R7 Beef Marketing). Being that resource scarcity is the only balancing loop of the system, the local and regional water availability and water scarcity determine the potential level of sustainability that a given area can reach (due to its internal carrying capacity). Using the Life Cycle Assessment approach, Ridoutt and Pfister (38) developed an adjustment factor that, in addition to the livestock water footprint estimate, allows the water footprint to be scaled to local conditions concerning a water scarcity index (Water Stress Index). The water stress index considers the local availability of the water resource, indicating the quantity of water used that is potentially removed from other activities. The application of this index means that the same production process could have a greater impact if carried out in conditions of scarce water availability in respect to areas with water abundance. Aside from regional considerations, the WF_B_ sustainability and consumer willingness to purchase beef include the slaughter and meat fabrication, retail, and household segments which were not included in this study. This proposed dynamic structure should be extended to these beef cattle segments to complete the full span of beef cattle water use and further identify specific leverage points for improvements in sustainability, WF_B_ levels, and more explicit determination of water uses. Thus, the question arises from this WF_B_ dynamic framework of how to capture specific water uses of the WF_B_ from cattle producers, meat processors, retailers, and consumers to understand meaningful long-term feedback relationships, delays, and the potential consequences.

## Conclusions

In conclusion, improved WF_B_ assessment is essential to achieve long-term improvements in livestock water use within and across the beef cattle supply chain. This study developed a dynamic framework to advance current WFA methods. The preliminary behavioral and sensitivity evaluations indicated that the framework is suitable to formulate the water footprint model for Texas with critical ruminant nutrition and growth equations using dynamic modeling software. A dynamic daily WF_B_ is likely to begin to resolve issues amongst existing WFA methodologies as it more accurately represents the dynamic nature of daily and total livestock water use. The CLD and their descriptions are essential and necessary to understand the complexity of the underlying structure and dominant loops that drive the long-term behavior of this system. Overall, freshwater challenges in agriculture livestock systems may be resolved by using this preliminary TXWFB framework to enhance the current livestock water footprint and supply chain assessment methods and quantify regional beef sustainability. Moreover, this study provides a new perspective for understanding the necessity for improved dialogue about WF_B_ sustainability within and across the beef cattle industry.

## Acknowledgments

We want to acknowledge the partial financial support by the 2017 Mini-Grant Opportunities to Enhance Teaching, Research, and Extension Capacity in the Department of Animal Science, and by the NIFA Hatch funding #TEX09123 Development of Mathematical Nutrition Models to Assist with Smart Farming Sustainable Production.

